# PAK1 activation drives divergent resistance mechanisms to aromatase inhibition and Tamoxifen in a luminal A breast cancer model

**DOI:** 10.1101/2025.04.28.651009

**Authors:** Luisa Schwarzmüller, Janina Müller, Efstathios-Iason Vlachavas, Lukas Beumers, Lena Fleischhacker, Sara Burmester, Dominic Helm, Cindy Körner, Stefan Wiemann

## Abstract

Breast cancer is the most frequent malignancy and the second leading cause of cancer-related mortality in women worldwide, and is a heterogeneous disease regarding driver mechanisms, therapies, and outcome. Estrogen receptor-positive (ER+) tumors are treated with endocrine therapies such as Tamoxifen or aromatase inhibitors (AI), aimed at disrupting tumor-promoting estrogen signaling. While these therapies are initially effective, resident tumor cells can develop resistance, leading to relapse. The p21-activated kinase 1 (PAK1) is a key regulator of multiple oncogenic signaling pathways and has been implicated in resistance to Tamoxifen. However, it remains unclear, whether PAK1 also affects the response to other endocrine therapies.

Using global phosphoproteomic profiling of endocrine resistant cell line models, we observed elevated PAK1 activity in cells resistant to Tamoxifen and to estrogen deprivation, respectively, with further increases following EGF stimulation. Pharmacological inhibition of PAK1 effectively reduced cell proliferation in both resistance models, however, with distinct effects on PAK1 downstream substrates and pathways. In the Tamoxifen resistance context, PAK1 reduces apoptosis via inhibition of the pro-apoptotic protein BAD, whereas in the AI resistance model PAK1 rather triggers proliferation via RAF1. Our findings position PAK1 as a common mediator of resistance to different endocrine therapies and offer insights into the distinct signaling networks that are involved.

In summary, our results suggest that targeting PAK1 may present a novel therapeutic strategy to overcome endocrine therapy resistance in ER+ breast cancer, potentially enhancing the efficacy of both Tamoxifen and AI treatments.

## Introduction

Breast cancer has remained the most frequent malignancy and the second leading cause of cancer-related death in women worldwide^1,2^. As breast cancer comprises a heterogeneous group of diseases, molecular subtyping of tumors is commonly performed in the clinics. This is commonly based on the assessment of estrogen receptor (ER), progesterone receptor (PR), human epidermal growth factor receptor 2 (HER2), and Ki-67 expression^3^. Around two-thirds of breast tumors are classified as ER+^4^ and the respective patients are treated with endocrine therapies to interfere with estrogen signaling^4,5^. Tamoxifen, a modulator of receptor activation, and aromatase inhibitors depriving tumors of estrogen, are first-line therapies in pre-and postmenopausal patients, respectively^5^. Despite initial effectiveness of endocrine therapies, up to 40% of patients presenting with late-stage disease at diagnosis ultimately develop resistance, relapse, and have a mostly poor prognosis^6^. Resistance to endocrine therapies can be mediated by various mechanisms, including mutations in *ESR1*^7^, the gene encoding ERα, upregulation of receptor tyrosine kinases, notably EGFR and HER2, and activation of downstream MAPK and PI3K pathways^8–14^. Several of these resistance mechanisms of endocrine resistance have been uncovered with help of therapy-resistant cell line models, which are commonly generated by long-term cultivation of cancer cell lines under therapeutic selective pressure^15^. Along these lines, we have previously described the upregulation of EGFR and HER2 signaling in two MCF7 cell line models of endocrine resistance^16^. These models had been generated by long-term cultivation of MCF7 cells in presence of tamoxifen (TAMR) or in media deprived of estrogen (LTED), the latter being a model for clinical aromatase inhibition.

PAK1, the p21-activated kinase 1, has been suggested as one possible player in endocrine therapy resistance, specifically in the context of tamoxifen treatment^17,18^. PAK1 is a central node that integrates signals from CDC42/RAC1, PI3K/AKT, EGFR/HER2/MAPK, and Wnt/β-Catenin pathways to regulate diverse cellular functions, including cytoskeletal organization, cell growth, survival, and migration^19–21^. Activated PAK1 signaling has been implicated in multiple cancer types and several small molecule inhibitors have been developed^22,23^. In breast cancer, the *PAK1* gene is frequently amplified, especially in the ER+ subtype^24,25^. Holm *et al.* found high expression of *PAK1* in 19% of premenopausal ER+ breast cancer patients. This was strongly correlated with higher malignancy and the patients did not respond to Tamoxifen treatment, evaluated by recurrence-free survival^17^. Gonzalez et al. showed that *PAK1* phosphorylation increased two-fold in response to tamoxifen treatment for 24 hours. Additionally, they showed that in tamoxifen-resistant MCF7, inhibition of the RAC1-PAK1 signaling axis using a RAC1-inhibitor, restored the anti-proliferative effects of tamoxifen^18^. While the role of PAK1 in therapy-resistance to tamoxifen thus seems to be established, it remains unknown whether PAK1 might have a causal relation also with resistance to estrogen deprivation via inhibition of aromatase. Such activity might prioritize inhibition of PAK1 as a potential clinical strategy to treat endocrine therapy-resistant tumors in post-menopausal patients.

Since cellular signaling processes are highly dynamic and mainly regulated at the protein level via post-translational mechanisms, such as phosphorylation^26^, extending molecular analyses beyond genomic and transcriptomic data is crucial to also assess the functional makeup of cancer cells. Here, we employed global, quantitative mass spectrometry (MS)-based total and phosphoproteomics to gain a detailed understanding of two endocrine therapy resistance models, MCF7 LTED and TAMR, representing resistance to aromatase inhibition and tamoxifen, respectively. The use of global phosphoproteomics data allowed us to assess the activities of kinases, which are frequently the drivers of cancer and are prime targets of therapeutic drugs^27^. In time-resolved phosphoproteomics experiments we observed PAK1 as one of the most active kinase in both LTED and TAMR compared to WT and saw further, prolonged PAK1 activation upon EGF stimulation. Both resistance models displayed similar dependence on PAK1 for proliferation. Pharmacologic inhibition of PAK1 identified common, however, also unique effects on PAK1-targets in LTED and TAMR conditions. While PAK1 drives proliferation in LTED, it rather inhibits apoptosis in TAMR. Our analyses revealed that PAK1 plays a central role in both LTED and TAMR resistance models and, by regulation of different downstream targets, might be a common target to interfere with endocrine therapy resistance.

## Results

### PAK1 is strongly overexpressed and activated in the MCF7 LTED cell line model

To study the complex mechanisms underlying endocrine therapy resistance in an isogenic background, we utilized two endocrine therapy-resistant cell line models that have been derived from the luminal A breast cancer cell line MCF7. The parental cell line had been cultivated either in the presence of tamoxifen or deprived of estrogen for 12 months, resulting in long-term estrogen-deprived (LTED) and tamoxifen-resistant (TAMR) cell lines, respectively^15,16,28,29^. The latter condition mimics resistance to therapeutic inhibition of aromatase.

Using the MCF7 LTED and TAMR models, we performed unbiased full and phosphoproteome analysis employing data-independent acquisition mass spectrometry (DIA MS), to assess changes in protein levels and pathway activities upon acquisition of endocrine therapy resistance. Full proteome analysis identified a total of 7,876 protein groups with at least 95% overlap between biological replicates (Suppl. Figure 1A, B). Principal component analysis suggested distinct adaptations in the two resistant cell line models (Figure 1A) and Reactome pathway overrepresentation analysis revealed several uniquely deregulated pathways, such as upregulated cholesterol biosynthesis in LTED and higher abundance of proteins involved in gluconeogenesis in TAMR, suggesting treatment-specific metabolic alterations (Figure 1B, Supplementary Figure 1C, D). In line with our proteomic findings, increased cholesterol synthesis has previously been implicated with endocrine therapy-resistance specifically in the LTED-condition by Nguyen et al. who reported that 27-hydroxycholesterol, a downstream metabolite of cholesterol, can activate ERα and thus compensate for the lack of estrogen^15^.

**Figure 1:**
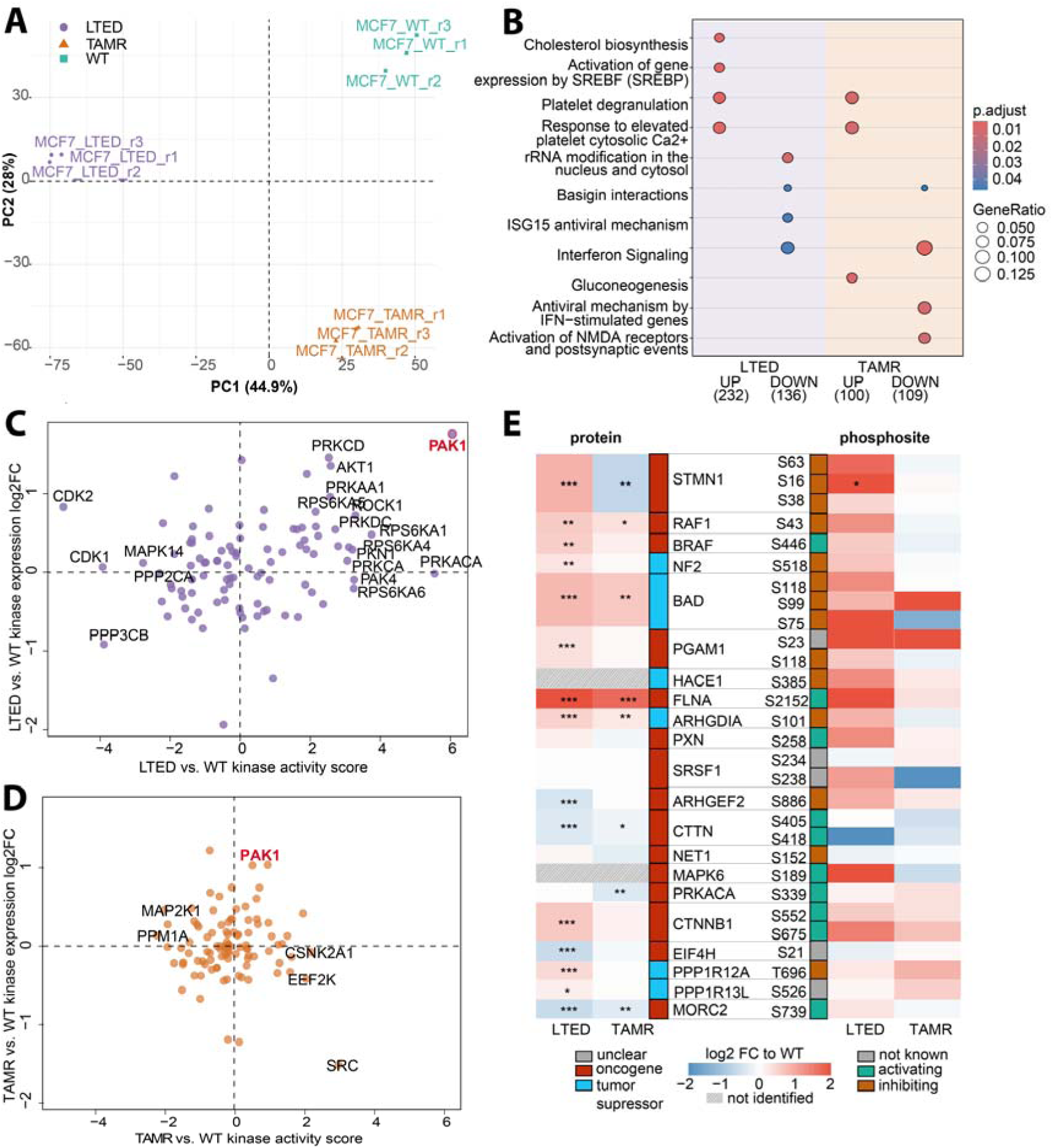
Proteomic and phosphoproteomic changes in MCF7 LTED and TAMR compared to WT cell lines. Cells were lysed in RIPA buffer, followed by 30 minutes of DNase/Benzonase treatment on ice. Protein clean-up and digestion was performed using manual SP3 with 500 µg protein input. Peptides were enriched for phosphorylation and both the full and phosphoproteome fractions were analyzed by LC-MS/MS for 120 minutes in DIA mode. Peptides and proteins were identified and quantified using Spectronaut (v17) in directDIA+ mode. Phosphorylated peptides were collapsed to the site-level using the Perseus plug-in PeptideCollapse with a localization cut-off at 0.95. (A) Principal component analysis of the full proteome analysis for LTED, TAMR, and WT conditions (n=3 for each condition). (B) Statistical analysis was performed using the eBayes function of the limma R package. Reactome pathway overrepresentation analysis was performed, using a threshold of fold-change > 2 (UP) or < 0.5 (DOWN) and an adjusted p-value < 0.05. (C, D) Statistical comparison was performed on the identified, localized and quantified phosphosites. Kinase activities were calculated on the t-statistics of the respective comparison, using decoupleR and Omnipath^35^. Kinase activities and fold-changes of protein abundance levels compared to MCF7 WT were plotted for LTED (C) and TAMR (D). Kinases with activity scores greater than 2 or below-2 are highlighted. PAK1 is additionally highlighted in TAMR. (E) Abundance of PAK1 target proteins and phosphorylation levels of known PAK1 substrate sites on these proteins are shown as log2 fold-change compared to WT. Significant p-values after BH-adjustment are highlighted: *** = p-value < 0.001, ** = p-value < 0.01, * = p-value < 0.05.

Signaling pathways are not only regulated at the level of protein abundance but more frequently by post-translational modifications, mostly phosphorylation, of central proteins^26^. To reveal potential mechanisms and druggable drivers of therapy resistance, we utilized the quantitative phosphoproteome data in combination with the Omnipath^30^ database of known kinase-substrate relationships to determine kinase activities. These were calculated based on T-statistics, in comparison to WT, of the intensities of known phosphosite substrates of each kinase. Here, PAK1 stood out as the most significantly activated kinase and was also the protein with the highest upregulated protein expression level in the LTED cells (Figure 1C). In contrast, PAK1 activity was not increased in TAMR cells (Figure 1D). Since PAK1 had already been described as a driver in acquired tamoxifen-resistance^17,18^, we continued investigating the context-specific functions PAK1 might have in the LTED context. There, the protein abundance of known downstream PAK1 target proteins with established oncogenic activities was upregulated, such as RAF1^31^, BRAF^32^, CTNNB1^33^, and FLNA^34^ (Figure 1E).

Next, we inspected the phosphorylation status of these proteins at their respective PAK1 substrate sites annotated in the Omnipath database. This analysis confirmed preferential phosphorylation of activating marks in PAK1 target proteins encoded by oncogenes (e.g., BRAF, FLNA, CTNNB1) and inactivating marks in tumor suppressor genes (e.g., NF2, BAD, PPP1R12A) (Figure 1E).

In conclusion, total and phosphoproteome analysis of the two resistance models revealed context-specific adaptations compared to the parental MCF7, e.g. in proteins involved in cholesterol biosynthesis or gluconeogenesis in LTED and TAMR, respectively. Combining the proteomic and phosphoproteomic data suggested that PAK1 might be a candidate kinase contributing to therapy resistance, particularly in the LTED context.

### Inhibition of PAK1 impairs viability of both endocrine therapy resistance models

As PAK1 is a known driver in acquired tamoxifen-resistance^17,18,36^ and since we had seen high levels and strong activation of PAK1 particularly in our MCF7 LTED model, we hypothesized that PAK1 might be involved also in resistance to estrogen deprivation by aromatase inhibitors. To test whether PAK1 activity is relevant for survival and proliferation in our resistance models, we treated the WT, LTED and TAMR cell lines with different doses of PAK1-specific small molecule inhibitor NVS-PAK1-1^37^ and quantified the effects on cell proliferation. Indeed, the LTED (IC50: 5.4 µM) and the TAMR (IC50: 7.2 µM) cell lines were significantly more sensitive to PAK1 inhibition than the WT cells (IC50:15.7 µM, Figure 2A) after seven days of treatment. This suggests that proliferation of both endocrine therapy resistance models relies on active PAK1 signaling. We thus hypothesized that the dependence on PAK1 might be a common mechanism beyond tamoxifen, to compensate for the loss of estrogen-stimulated signaling. To test this for the LTED model, we next re-introduced estrogen into the culture medium of LTED cells and measured *PAK1* mRNA levels after 4, 8 and 12 weeks. Upon re-exposure to estrogen, *PAK1* expression strongly decreased in the LTED cells and remained low for the entire duration of the experiment (Figure 2B), suggesting a direct relation between estrogen availability and PAK1 levels.

**Figure 2:**
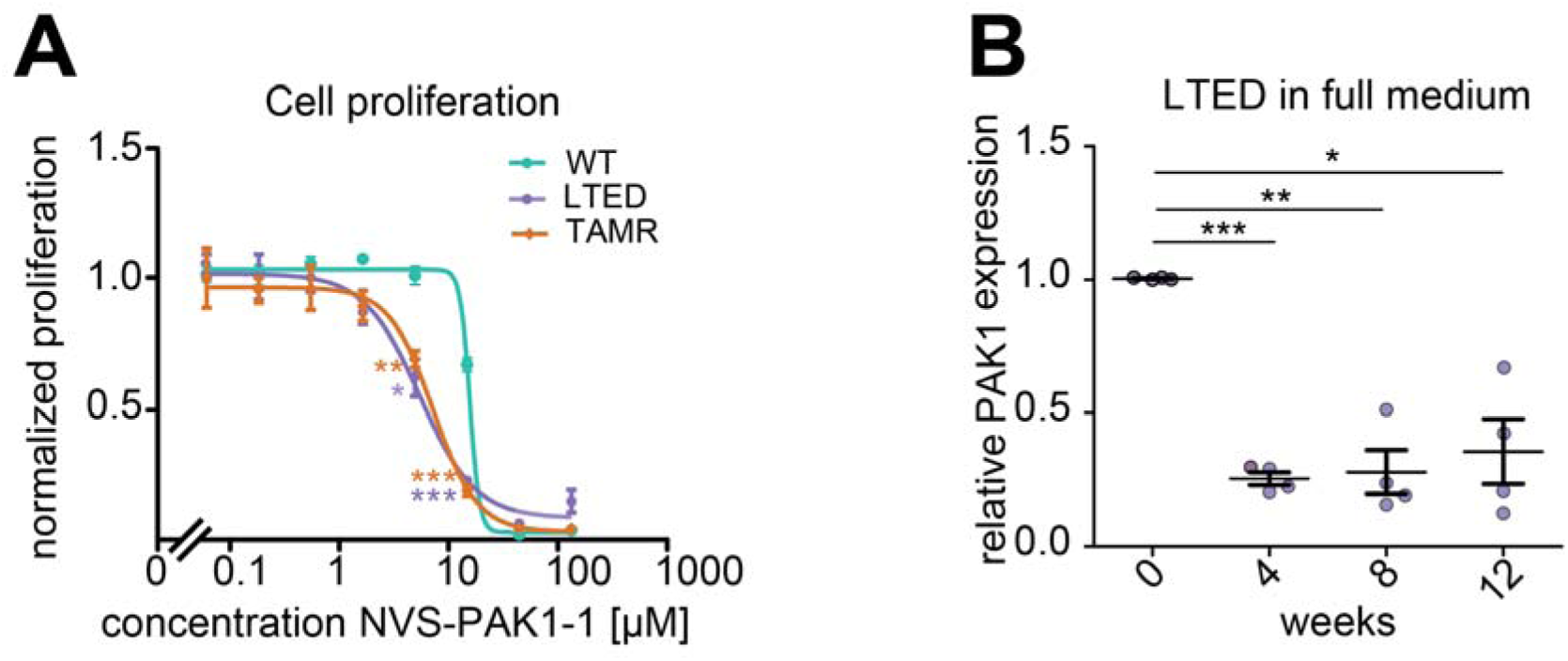
PAK1 inhibitor treatment and PAK1 expression upon re-exposure to estrogen. (A) Cells were plated in 96-well plates and treated with different concentrations of PAK1 inhibitor NVS-PAK1-1 (Cayman Chemical, MI, USA) or DMSO control. Proliferation was determined by counting the numbers of nuclei in alive cells at day 7 and related to respective numbers at day1, relative to the DMSO control. n=3 biological replicates, 2-sided T-test, * = p<0.05, ** = p<0.01, *** = p<0.001. (B) Estrogen was re-introduced into LTED medium for 4, 8 or 12 weeks and *PAK1* expression was measured via RT-qPCR. Expression is shown relative to medium without estrogen. n=4 biological replicates, 2-sided T-test, * = p<0.05, ** = p<0.01, *** = p<0.001.

Here, we have shown that both endocrine therapy-resistant cell models were more sensitive to PAK1-inhibition compared to WT cells, suggesting that PAK1 is required for proliferation and survival when estrogen signaling is abrogated by either depletion of estrogen (i.e. aromatase inhibition) or by selective modulation of estrogen receptor signaling (i.e. tamoxifen treatment).

### EGF stimulation triggers prolonged activation of downstream pathways and further enhanced PAK1 activity in LTED and TAMR cell line models

In a previous study, we had shown that MCF7 expresses moderate levels of EGFR and responds to EGF stimulation with a fast and transient activation mostly of RAS-RAF-ERK signaling^38^. Furthermore, an activating mutation in PIK3CA that is present in MCF7 triggers a prominent basal activation also of the PI3K-AKT-mTOR pathway^38^. In addition to MAPK and AKT signaling, EGFR has been described to activate PAK1 via RAC1 or NCK^39,40^. As regulation of PAK1 via RAC1 has been associated with tamoxifen resistance^36^, we hypothesized that EGF stimulation might modulate PAK1 activity also in our endocrine resistance models. To test this, we performed a time-resolved experiment with EGF stimulation to assess PAK1 signaling and pathway activation over time, again using an MS-based full and phosphoproteomic readout.

The activation of pathways downstream of EGFR is commonly estimated based on the phosphorylation levels of classical regulatory sites in selected proteins involved in EGFR-signaling, such as MAPK3/ERK1 T202/Y204 and MAPK1/ERK2 T185/Y187. Indeed, the phosphorylation levels at these sites were strongly upregulated upon EGF stimulation as compared to growth factor-deprived cells (Suppl. Figure 2). Accordingly, we observed increased activities particularly of kinases in the MAPK/AKT/mTOR/S6K signaling pathways downstream of EGFR in WT, LTED as well as TAMR cell lines in response to EGF stimulation (Figure 3A).

**Figure 3:**
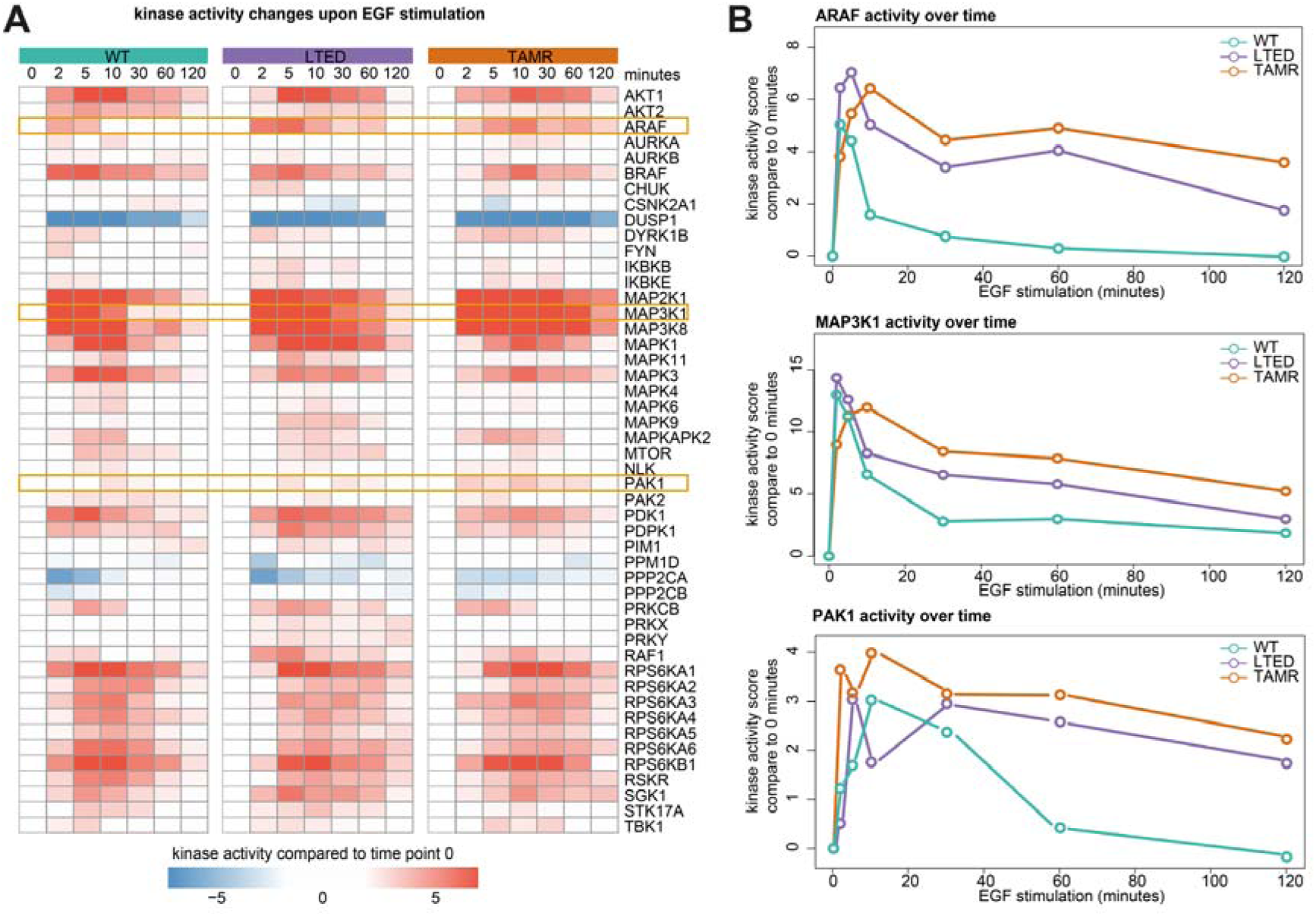
Phosphoproteomic changes in MCF7 cell lines upon EGF stimulation. Cells were cultivated without growth factors for 24 hours, followed by stimulation with 5 nM EGF. At indicated time points, cells were lysed in RIPA buffer, followed by 30 minutes of DNase/Benzonase treatment on ice. Protein clean-up and digestion was performed using manual SP3 with 500 µg protein input. Peptides were enriched for phosphorylation and both the full and phospho proteome fractions were analyzed by LC-MS/MS for 120 minutes in DIA mode. Peptides and proteins were identified and quantified using Spectronaut (v17) in directDIA+ mode. Phosphorylated peptides were collapsed to the site-level using the Perseus plug-in PeptideCollapse with a localization cut-off at 0.95. Statistical comparison was performed using the eBayes function of limma (v. 3.58.1). Kinase activities were calculated on the t-statistics of the respective comparison, using decoupleR and Omnipath. (A) Kinase activities within each cell line compared to unstimulated condition (timepoint 0) (B) Activities of PAK1, RAF1 and MAP3K1 over time.

We next assessed the dynamics of kinase activities in the two endocrine resistance models as compared to the parental WT cell line. While the global response of kinases to EGF stimulation appeared to be similar in the therapy resistance models compared to MCF7 WT (Figure 3A), we utilized the time-resolved analysis to reveal differences in the activation-dynamics of phosphosites and their respective kinases. We applied two-way ANOVA and identified 279 phosphosites with significantly different dynamics upon EGF stimulation when comparing the resistant and WT cell lines (Suppl. Figure 3A). Most of the sites showed prolonged phosphorylation in LTED and TAMR with increased intensities lasting for 60 or even 120 minutes post EGF stimulation, while in the WT cell line most of these phosphosites returned back to baseline intensities within 30 minutes (Suppl. Figure 3A). Reactome pathway analysis revealed that these differentially phosphorylated sites prominently mapped to proteins related to RTK and MAPK signaling (Suppl. Figure 3B), indicating potentially prolonged RTK and MAPK signaling. Sixty of these phosphosites had an upstream kinase annotated in the Omnipath database. Specifically, ARAF, BRAF, MAP2K1, MAPK3, MAPK1, PDK1, AKT1, RPS6KB1, PRS6KA3, and RPS6KA1 were overrepresented as upstream kinases for this set of phosphosites with significantly altered time-resolved dynamics in the resistance models, indicating prolonged activation dynamics (Suppl. Figure 4).

Furthermore, PAK1 remained at higher activation levels in both LTED and TAMR even 120 minutes after EGF stimulation, while its activity in WT cells was back to the basal level after 60 minutes. Similarly, the activation of ARAF and MAP3K1 was more transient specifically in the WT cells (Figure 3B). The consistent finding of prolonged protein phosphorylation as well as prolonged kinase activation suggests that likely reduced activities of negative feedback loops ensure prolonged signal activities in the endocrine therapy-resistant cell lines, potentially contributing to altered phenotypes.

The consistently deregulated kinases, compared to WT, largely overlapped in LTED and TAMR across all time points of EGF stimulation. These included more active PAK1, BRAF as well as several members of the RPS6K family (Figure 4A). The phosphatase DUSP1 is less active in both resistance models compared to WT, especially at the later time points (Figure 4A), which might explain the prolonged signaling duration we observed^41^.

**Figure 4:**
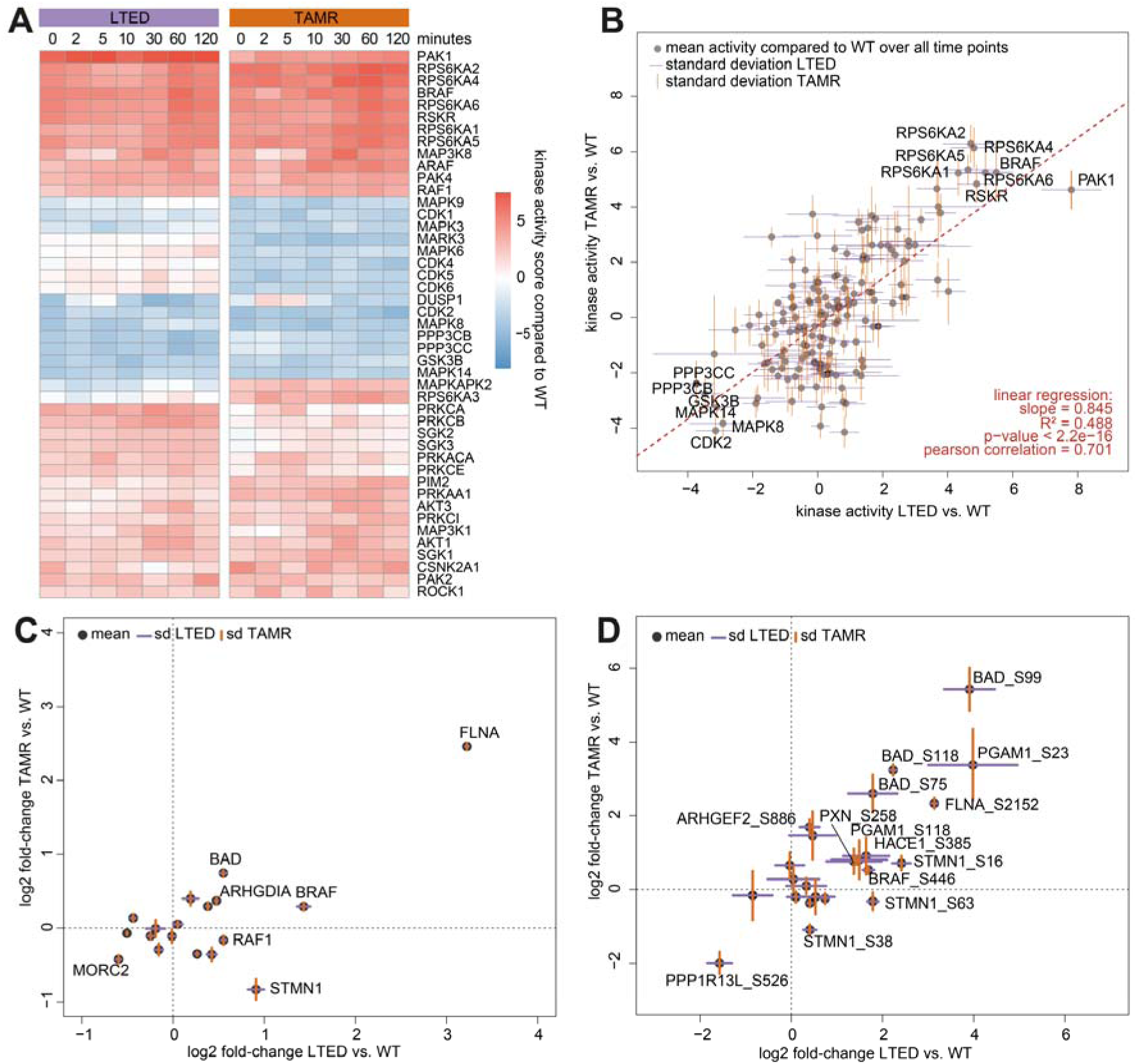
Kinase activity changes in LTED and TAMR compared to WT over the time course of EGF stimulation. (A) Kinases with absolute activity scores > 2 in LTED and TAMR compared to WT at each time point of EGF stimulation. (B) Mean and standard deviation of all kinases compared to WT over all time points. (C) Log2 fold-changes of full proteome protein abundance and phosphosite intensities of known PAK1 substrates in LTED compared to WT. Shown are mean ± SEM over all time points. Proteins with an absolute log2 fold-change > 0.5 and a significant difference to WT are labelled. (D) Log2 fold-changes of protein abundance and phosphosite intensities of known PAK1 substrates in LTED vs. WT and TAMR compared to WT. Shown are mean ± SD of all time points. Phosphosites with an absolute log2 fold-change > 1 and a significant difference to WT are labelled.

The degree of kinase activity deregulation was highly similar between LTED and TAMR with a Pearson correlation coefficient of 0.701 (Figure 4B). In contrast, kinase activities and their respective protein abundance were largely unrelated (Suppl. Figure 5), highlighting the importance of assessing protein or pathway activities rather than gene or protein expression only.

Again, PAK1 was found to be the most activated kinase in LTED compared to WT (Figure 4B). Interestingly, while PAK1 was only slightly more active in TAMR under normal growth conditions (score = 0.52, see Figure 1D), it was four times more active in the growth factor-deprived setting (Figure 4B). PAK1 is known to be activated by EGFR signaling especially under nutrient stress^39^. While TAMR cells clearly depend on PAK1 for proliferation also in normal growth medium (Figure 2A), serum starvation might be required for full activation of PAK1.

To zoom in on potential mechanisms downstream of PAK1 that might explain estrogen signaling-independent survival and proliferation, we next focused on the phosphorylation levels of known PAK1-substrates, as annotated in the Omnipath database. Highly elevated protein-levels of Filamin A (FLNA) were detected in both LTED and TAMR cells (Figure 4C) and the protein was also more strongly phosphorylated, specifically at serine residue S2152 (Figure 4D). Phosphorylation at this site by PAK1 has been described as part of a positive feedback loop where the interaction between FLNA and PAK1 enhances PAK1 kinase activity^42^. Also in line with our findings, FLNA has been associated with various cellular processes, including cell adhesion, proliferation, and DNA repair^42^ and has previously been implicated as a driver in chemotherapy resistance by activating MEK-ERK signaling^43^. The pro-apoptotic protein BCL2 Associated Agonist of Cell Death (BAD) was significantly more phosphorylated at residues S99, S118 and S75 in endocrine resistant cells. Especially in the TAMR condition, a more than 30-fold higher intensity of S99-phosphorylated BAD was measured compared to WT (Figure 4D). All three phosphorylation sites have been reported to inhibit the pro-apoptotic function of BAD^44^, suggesting a potential involvement in the estrogen-independent survival of the endocrine therapy-resistant cells.

In summary, the kinase activity landscapes we uncovered in response to EGF stimulation were mostly similar in LTED and TAMR models. PAK1 was consistently among the most strongly activated kinases in both resistance models, its activity further increased upon EGF stimulation, and demonstrated prolonged activation compared to WT cells. The dependence of endocrine therapy resistant cells on PAK1 signaling for survival and proliferation might also be related to increased phosphorylation of downstream substrates, like FLNA and BAD.

### Inhibition of PAK1 indicates distinct wiring of downstream pathways in the two resistance models

To further elucidate the signaling network downstream of PAK1 in both resistance models and to uncover how it might trigger estrogen-independent survival and proliferation, we acquired phosphoproteome data after short-term PAK1 inhibition. LTED and TAMR cells were treated with 6 µM of the PAK1-specific inhibitor NVS-PAK1-1, which is close to the previously determined IC50 concentrations for LTED and TAMR cells (see Figure 2). As expected, no significant changes were detectable in the full proteomes as compared to the DMSO control after one hour of PAK1 inhibition (Suppl. Figure 6A). In contrast, strong effects were visible in the phosphoproteomic data. Consistently, decreased activities of the kinases PAK1, CDK17, and most prominently, PAK2 were observed in both LTED and TAMR upon short-term PAK1 inhibition (Figure 5A). While the strong repression of PAK2 could potentially hint at an off-target effect of the PAK1 inhibitor, inspection of residues specifically phosphorylated by PAK2 revealed that its predicted lower activity was mainly based on the decreased phosphorylation at the auto-phosphorylation site S141 of PAK2 (Suppl. Figure 6B). Furthermore, the NVS-PAK1-1 inhibitor has been reported as a selective PAK1 inhibitor having an almost 60-fold higher selectivity for PAK1 over PAK2^45^. This might support a direct interaction between PAK1 and PAK2, which has indeed been described before^46^, rather than an off-target effect of the inhibitor. AKT1, MAPK1, MAPK3 and MAPK8 showed consistently increased activities upon PAK1 inhibition in both cell models, indicating a potential feedback loop between PAK1 and the AKT/MAPK pathways.

**Figure 5:**
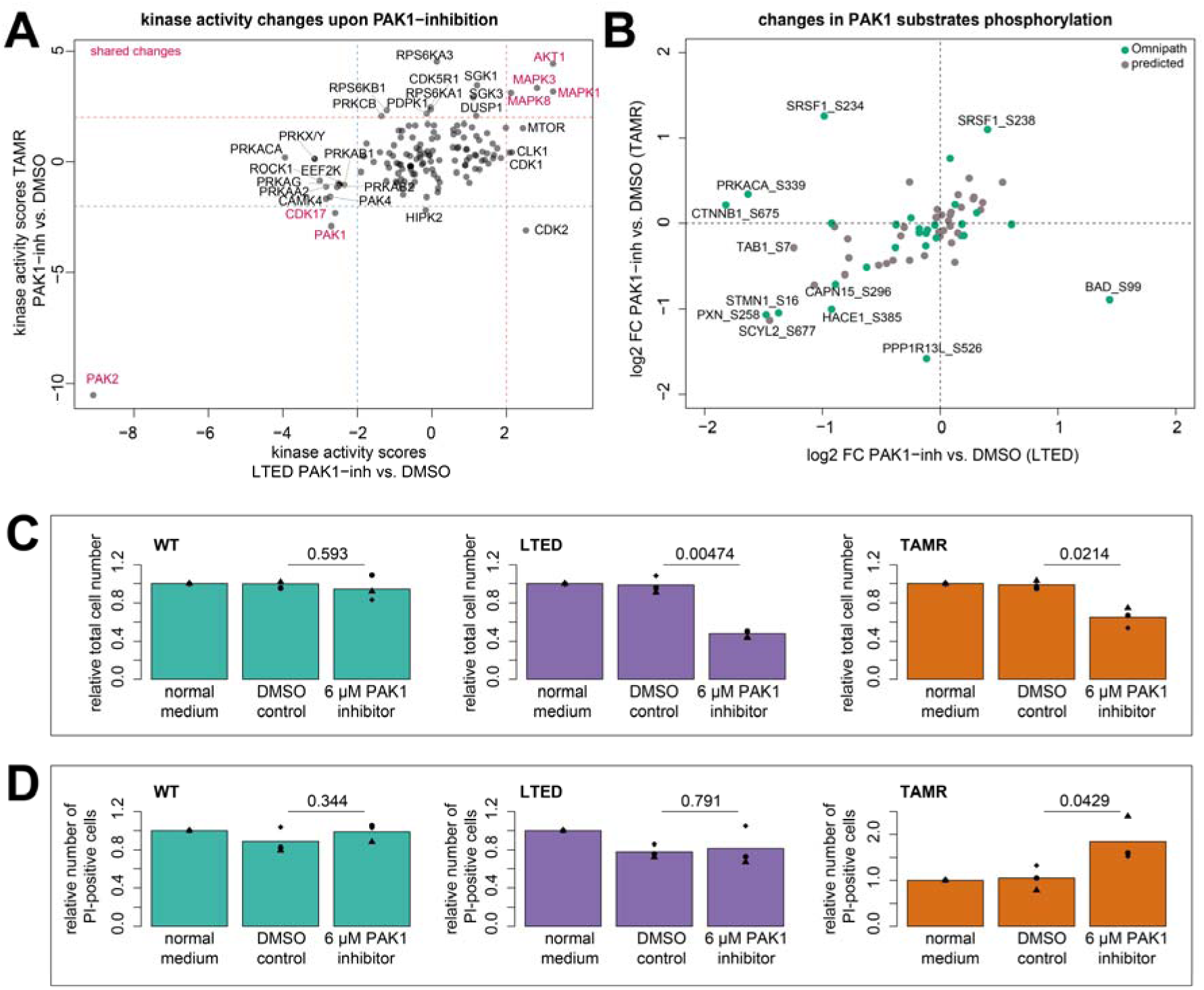
Immediate downstream effects of PAK1-inhibition in LTED and TAMR cells. MCF7 cell lines were treated with 6 µM PAK1 inhibitor for 1 hour and subsequently lysed in RIPA buffer. Proteins were prepared for LC-MS/MS analysis by automated SP3 and phospho-enrichment of 55 µg protein as input material. Both the full and phospho proteome fractions were analyzed by LC-MS/MS for 120 minutes in DIA mode. Peptides and proteins were identified and quantified using Spectronaut (v17) in directDIA+ mode. Phosphorylated peptides were collapsed to the site-level using the Perseus plug-in PeptideCollapse with a localization cut-off at 0.95. Statistical comparison was performed using the eBayes function of limma (v. 3.58.1). Kinase activities were calculated on the t-statistics of the respective comparison, using decoupleR and Omnipath. (A) Kinase activity scores upon PAK1 inhibition for 1 hour in LTED and TAMR compared to the DMSO control. (B) Known PAK1 substrates were retrieved from the Omnipath database, predicted PAK1 substrates were obtained from Johnson et al.^47^ by filtering for phosphosites with PAK1 in at least the 98^th^ percentile and ranking higher than 5. Log2 fold-changes of known and predicted PAK1 substrates’ intensities in the PAK1-inhibited condition compared to the DMSO control in LTED and TAMR, respectively. (C, D) Cells were treated with 6 µM PAK1 inhibitor for 7 days. Proliferation was measured by Hoechst staining. Apoptotic cells were detected with PI staining. The number of cells was compared to day1 and further normalized by the normal growth medium. Statistics was performed using a one-sided unpaired t-test to test for fewer living cells or more dead cells, respectively.

The effect of PAK1-inhibition further revealed that several PAK1 substrates annotated in the Omnipath database, specifically PXN (S258), NF2 (S518), and STMN1 (S16, S63) strongly decreased in both LTED and TAMR (Figure 4B), suggesting their direct regulation by PAK1 in both resistance models. Databases of known kinase-substrate networks, such as Omnipath, rely on experimentally validated relationships and might therefore miss out on so far unknown interactions. To identify new potential downstream substrates of PAK1, we made use of the target site prediction tool published by Johnson et al.^47^. Several of the PAK1 substrates predicted there, such as TAB1 (S7), CAPN15 (S296), and SCYL2 (S677) indeed showed decreased phosphorylation in response to PAK1 inhibition (Figure 4B), which supports them as putative PAK1 substrates, at least in the LTED and TAMR context.

Further, PAK1 inhibition exerted differential effects in LTED vs. TAMR, such as a decrease in phosphorylation of RAF1 (S43), PRKACA (S339), and CTNNB1 (S675) in LTED only, suggesting that PAK1 potentially acts via activating these known oncogenic driver proteins specifically in the LTED context. In contrast, inhibition of PAK1 strongly decreased phosphorylation of BAD at S99 exclusively in the TAMR-cells (Figure 4B). In line with a previous study which had associated this phosphosite with tamoxifen resistance^48^, our data suggests that in TAMR cells PAK1 might act through inhibition of BAD to promote estrogen-independent survival and proliferation.

To test this hypothesis, we treated MCF7 WT, LTED and TAMR cells with 6 µM PAK1-inhibitor and measured the total number of cells, as well as the number of dead cells after 7 days. Consistent with the data shown in Figure 2A, the proliferation of WT cells remained unaffected by PAK1 inhibition while, as expected, proliferation was reduced by 40-60% in both, LTED and TAMR cells (Figure 5C). However, the number of dead, PI-positive, cells was significantly increased only in the TAMR condition whereas cell death seemed to be unaffected by PAK1 inhibition in LTED cells as well as in the DMSO control (Figure 5D). This finding suggests that PAK1 acts via inhibiting apoptosis, potentially through BAD, specifically in the tamoxifen-resistance context.

## Discussion

Endocrine therapy resistance mechanisms in estrogen receptor α (ERα) positive breast cancer have been studied for several decades^49^. Multiple studies have elucidated potential resistance mechanisms, including estrogen-independent activation of ERα and the activation of alternative survival pathways, such as EGFR, HER2, MAPK and PI3K/AKT^11,16,50–54^. The development of endocrine therapy-resistant cell line models has significantly advanced those discoveries by enabling in-depth *in vitro* analysis of the molecular changes associated with resistance^50^. For instance, using independently established cell line models, Zhou et al. performed proteomic characterization of MCF7 TAMR cells and observed down-regulated ER signaling alongside the activation of alternative survival pathways^11^. Collectively, resistance mechanisms appear to manifold and might also derive from stochastic clonality during the outgrowth of resistant cells^55,56^.

In our study, we identified several proteomic and phosphoproteomic adaptations that potentially contribute to endocrine therapy resistance. Proteomic changes revealed notable differences in metabolic pathways such as cholesterol biosynthesis und gluconeogenesis. Cholesterol biosynthesis has been implicated in breast cancer initiation and progression, as reviewed by Kim et al.^57^. A preclinical study has demonstrated that combining tamoxifen treatment with cholesterol depletion (acetyl plumbagin) reduced cell proliferation in MCF7 and MCF7-LTED cells^58^. Furthermore, 27-hydroxycholesterol has previously been found to be a substitute for estrogen, in the same estrogen-deprived resistance model^15^ that we employed in our study.

At the phosphoproteome level, PAK1 emerged as a central kinase, showing strongly increased abundance and elevated activity prominently in LTED compared to WT cells. While the role of PAK1 as a resistance driver in TAMR has been previously shown indirectly via RAC1-inhibition^18^, we here confirmed its relevance by direct inhibition of PAK1. According to the literature, PAK1 has been associated with endocrine resistance only when the *PAK1* gene was strongly overexpressed^17,21,24^. In line with these findings, we observed increased PAK1 activity in comparison to WT cells, and further increased PAK1 activation upon EGF stimulation. These alterations were indeed accompanied by elevated PAK1 levels in the LTED condition, however, not in TAMR cells. Thus, we would have missed PAK1 as a putative driver specifically in the TAMR cell line if this had been based only on total protein levels. These findings support the importance of assessing kinase activities rather than relying solely on protein abundance. This apparent disparity we observed between protein levels and activities (compare Suppl. Figure 5C, D) has been described before and has been associated with various mechanisms, such as post-translational modifications, co-factor requirements, or subcellular localization^59,60^.

PAK1 activity further increased upon stimulation with EGF and this was sustained for a longer time in LTED and TAMR compared to WT. Even minor changes in the duration of activity of these proteins can have substantial biological effects on cellular proliferation and survival dynamics^61^. Structural changes in the EGFR that are induced upon ligand-binding have been associated with differential duration of growth factor activity and different phenotypic effects these induce^62^. The duration of signaling pathways downstream of EGFR, like MAPK, is further regulated by several other mechanisms, including positive and negative feedback loops, the availability of scaffold proteins, and cross-talk between different MAPK pathways^61,63^. The extended activation of downstream signaling we observed in PAK1-activated cells is thus likely to contribute to differential phenotype alterations also in the cell line systems we investigated, including endocrine resistance. Additionally, we observed prolonged activation of MAP3K1 and ARAF specifically in the LTED and TAMR resistant cell lines further supporting a role of MAPK signaling in resistance. In summary we have shown that LTED and TAMR cell line models share a number of features, such as reliance on PAK1 for estrogen-independent proliferation.

While the integration of unbiased proteomic and phosphoproteomic analysis in cancer research is increasingly applied, its clinical application remains constrained by the limited understanding on the functional relevance of most phosphosites. Currently, only about 5% of identified phosphosites have known upstream kinases and well-characterized functions^64^. As a result, current phosphoproteome analyses is restricted to a few kinases and their substrates. New tools for the prediction of kinase/substrate and phosphosite/functionality interfaces promise to broaden application-spectrum of phosphoproteomics^47,64–67^. Here, we utilized the predictions provided by Johnson et al.^47^ to identify potential new PAK1 substrates. While several of these putative PAK1 substrates are shared with many other kinases, inhibitor experiments, like the one we applied in our study, will help to eventually validate more. One interesting candidate is SCYL2, a poorly studied protein involved in vesicle transport and associated with endosomes and the trans-Golgi network. Our data suggest that SCYL2, specifically at its S677 site, is a novel PAK1 substrate in the context of endocrine therapy resistant MCF7.

In conclusion, PAK1 activation seems to contribute to endocrine therapy resistance in both LTED and TAMR cell line models. The resistant MCF7 cells depend on PAK1 signaling for cell proliferation and are, therefore, sensitive to PAK1 inhibition. In-depth investigation of the signaling networks revealed common, but also distinct, downstream targets regulated by PAK1 in the two different isogenic resistance models indicating cellular plasticity. Further studies including preclinical models or investigation of the impact PAK1 might have on the inhibition of primary breast cancer organoids derived from patients having relapsed under endocrine therapy are required to pave the way towards potential targeting of PAK1 as a novel strategy in combinatorial therapy.

## ACKNOWLEDGEMENTS

We thank Ursula Klingmüller and Jeroen Krijgsveld for providing access to an MSCoreSys network SMART-CARE [031L0212B] funded Orbitrap Exploris 480 mass spectrometer.

We thank Luca Magnani for providing the LTED and TAMR MCF7 cell lines used in this study.

## AUTHOR CONTRIBUTIONS

L.S., J.M., L.B., D.H., C.K., and S.W. conceived the study. L.S., J.M., L.B., L.F., and S.B. designed and/or performed experiments. L.S., and E.V. analyzed and visualized the data. L.S., C.K., and S.W. wrote the manuscript. All authors read and approved the final manuscript.

## FUNDING

Open Access funding enabled and organized by Projekt DEAL.

## COMPETING INTERESTS

The authors declare no competing interests.

**ADDITIONAL INFORMATION**

**Supplementary information**

## Materials & Methods

### Cultivation of cell lines

MCF7 (Cellosaurus CVCL_0031) cell lines were a kind gift from Prof. Luca Magnani (Imperial College London)^15^. Cell lines were authenticated using STR profiling and regularly tested for potential mycoplasma contamination. Cell lines were cultivated in Dulbecco’s Modified Eagle Medium (DMEM) supplemented with 10% FCS, 50 units/mL penicillin, 50 µg/mL streptomycin sulfate (Invitrogen AG, Carlsbad, CA, USA), and 10−8 M 17-ß-estradiol (E2, Sigma-Aldrich, Saint-Louis, MI, USA) at 37°C with 5% CO2 in a humidified incubator. Tamoxifen resistant and long-term estrogen deprived cells were either maintained in presence of 100 nM of the active metabolite of tamoxifen, 4-hydroxytamoxifen (4-OHT, Sigma-Aldrich) or cultured in estrogen-free DMEM (w/o phenol red) supplemented with 10% charcoal stripped FCS, respectively.

### Inhibitor treatment and proliferation assay

Cells were plated in black clear-bottom 96-well plates (Greiner Bio-One International GmbH, Kremünster, Austria) and treatment with the PAK1 inhibitor NVS-PAK1-1 (Cayman Chemical, MI, USA) resuspended in dimethyl-sulfoxide (DMSO, Sigma-Aldrich) for seven days. Cells were stained with the fluorescent intercalating dye Hoechst 33342 and measured using the IXM XLS microscope (Molecular Devices, San Jose, CA, USA) and the Molecular Device Software. Proliferation was determined relative to DMSO control.

### EGF stimulation

Cells were starved with FCS-depleted media for 24 hours. The following day, cells were stimulated with 5 nM epidermal growth factor (EGF, Sigma-Aldrich, #E9644), and samples were collected at the indicated time points.

Following two washing steps with PBS, cells were lysed in RIPA buffer (Thermo Fisher Scientific) that was supplemented with 10 mM NaF, 1 mM Na3VO4, cOmplete EDTA-free protease inhibitor (Merck), PhosSTOP phosphatase inhibitor (Merck), 250 U/mL Benzonase (Merck), and 10 U/mL DNase (Qiagen). Lysates were kept on ice for 30 minutes and then centrifuged for 30 minutes at 15,000xg and 4°C. Supernatants were then transferred to fresh tubes. Protein concentrations were determined using the Pierce BCA assay (Thermo Fisher Scientific) and lysates were stored at-80°C until further use.

### Sample preparation workflow for phosphoproteomics

Five hundred µg of protein were precipitated according to Wessel and Flügge^68^, protein pellets were dried for 15 minutes at room temperature, and stored at-20°C.

For tryptic digestion, protein pellets were dissolved in 8 M urea, supplemented with 100 mM NaCl, 50 mM TEAB, cOmplete EDTA-free protease inhibitor (Merck) and PhosSTOP phosphatase inhibitor (Merck). Disulfide bridges were reduced with 10 mM DTT for 1 hour at 27°C, then alkylated with 30 mM IAA for 30 minutes at room temperature. Reactions were quenched with 10 mM DTT and incubation for 15 minutes at room temperature.

Lysyl endopeptidase (LysC, Fujifilm) was added in a 1:100 enzyme-to-protein ratio, and proteins were digested for 4 hours at 30°C. Then the urea concentration was diluted to 1.6 M, Trypsin was added in a 1:50 ratio and the digestion continued for 16 hours at 37°C. Formic acid (FA) was added to a final concentration of 2% (v/v) to terminate digestion.

Peptides were desalted using Sep-Pak C18 cartridges (Waters). The cartridges were conditioned with 100% acetonitrile, washed with 80% acetonitrile containing 0.6% acetic acid and equilibrated twice with 2.5% formic acid. Peptide solutions were loaded, and the flow-through was collected and loaded a second time. Bound peptides were washed four times with 2.5% formic acid and then eluted twice with 80% acetonitrile containing 0.6% acetic acid. The eluate was vacuum centrifuged to dryness and stored at-20°C.

For enrichment of phosphorylated peptides, an Immobilized Metal Affinity Chromatography (IMAC) column (Thermo Fisher Scientific) was charged with 25 mM iron chloride (FeCl_3_) in 100 mM acetic acid at a flow rate of 0.2 mL/min for 30 minutes. Subsequently, the column was rinsed with 0.1 % formic acid at a flow rate of 4 ml/min for 4 hours. Dried peptides were reconstituted in 30% acetonitrile with 0.07 % TFA, and were then loaded onto the IMAC column with a flow rate of 0.2 mL/min. Elution of phosphorylated peptides was achieved with a gradient of ammonium (NH_4_OH). The phospho fraction was collected and vacuum centrifuged to dryness.

Finally, the enriched phosphopeptides were desalted by using Stop and Go Extraction (STAGE) tips. Briefly, three layers of C18 material (Supelco) were placed in a pipette tip, pre-wetted with methanol, washed with 80% acetonitrile in 0.1% trifluoroacetic acid (TFA) and equilibrated with 0.1% TFA. The dried peptides were reconstituted in 0.1% TFA and loaded onto the STAGE tip. Subsequently, the peptides were washed with 0.1% TFA, followed by two elution steps with 60% and 80% acetonitrile, respectively. The eluate was vacuum centrifuged to dryness and stored at-20°C until further analysis.

For the MS analysis of the PAK1 inhibitor-treated samples, protein clean-up was performed with 55 µg protein material as input following the automated single-pot solid phase enhanced sample preparation (SP3) workflow adapted from Müller et al.^69^ on the Agilent Bravo liquid handling robot. Proteins were digested with Trypsin in a protease-to-protein ratio of 1:25 for 16 hours at 37°C. Peptides were vacuum centrifuged to dryness and subsequently enriched for phosphorylation on the Agilent Bravo liquid handling robot following the Agilent application note (https://www.agilent.com/cs/library/applications/5991-6073EN.pdf).

### Liquid Chromatography

For LC-MS/MS analysis, peptides were dissolved in ULC/MS grade water containing 0.1% trifluoracetic acid (TFA) and 2.5% 1,1,1,3,3,3-Hexafluoro-2-propanol (HFIP), followed by sonication for 5 minutes. For phospho-enriched samples, the reconstitution buffer additionally contained 50 mM citric acid. The samples were transferred to autosampler vials and placed in the autosampler module of the Ultimate 3000 (Thermo Fisher Scientific) liquid chromatography (LC) system. The LC was operated at a flow of 300 nL/min and the column was heated to 35°C. Peptides were first loaded onto a trapping column (Thermo Fisher Scientific, 160454) in the presence of 98% loading buffer A (0.1% TFA in water) and 2% loading buffer B (0.1% TFA in acetonitrile). Peptides were then transferred to the analytical column (Waters, 186008795, BEH C18 130Å 1.7 µm 75×250 mm), from which they were separated by hydrophobicity with a linear gradient of 4-30% acetonitrile. For phosphopeptide-enriched samples, a gradient of 2-28% acetonitrile was applied. Separated peptides were ionized by electrospray ionization (ESI) with an applied voltage of 2200 V.

### Mass Spectrometry measurement

The Orbitrap Exploris 480 mass spectrometer (Thermo Fisher Scientific) was operated in DIA mode. MS1 scans were acquired at a resolution of 120K and covered the range from 350 - 1400 m/z. Maximum injection time was 45 ms and the automated gain control (AGC) target was set to 3e6. MS2 scans were acquired in 47 precursor isolation windows of variable width and 1 m/z overlap that covered the range from 400 – 1000 m/z (up to 1200 m/z for phospho-enriched samples). The orbitrap was operated at a resolution of 30K and a normalized collision energy of 28% (26% for phospho-enriched samples) was applied. Maximum injection time was 54 ms and the automated gain control (AGC) target was set to 1e6. The total cycle time of the method equaled 3.6 seconds.

### MS raw data search

Peptide and protein identification and quantification from DIA raw data was performed with Spectronaut (Biognosys, version 17) in library-free (directDIA+) mode. For phosphopeptides, the newly developed identification and localization algorithm implemented in Spectronaut was used^70^. Identified phosphorylated peptides were site-collapsed using the Perseus^71^ (version 1.6.2.3) plug-in PeptideCollapse, as described by Bekker-Jensen et al.^70^, with a localization cut-off at 0.95.

### Data preprocessing and differential expression analysis

Data were further analyzed using R^72^ (v. 4.0.2) implemented in RStudio^73^ (v. 1.3.1093). Protein or phosphosite intensities were normalized using the R packages vsn^74^ (v. 3.70.0) and SummarizedExperiment (v. 1.32.0), accordingly. Moreover, we adapted a filtering method available in the PhosR package^75^ (v. 1.4.0), to keep proteins or phosphosites that are present in at least 2 out of 3 replicate samples in at least one condition. Differential protein expression analysis for the respective comparisons was performed using the limma^76^ R package (v. 3.50.3). Resulting p-values were adjusted for multiple hypothesis testing using Benjamini-Hochberg method.

### Inference of differentially activated kinases

Relative kinase activities were determined based on the T-statistics of the differential phosphosite abundance analysis. The OmnipathR^30^ package (v. 3.2.8) was used to retrieve kinase-substrate relationships. To ensure the reliability of our approach, we excluded interactions that were solely present in ProtMapper^77^ and not corroborated by any other source. Subsequently, the decoupleR^35^ package (v. 2.1.6) was employed to estimate the relative activity scores by weighting the molecular readouts of its targets according to their mode of regulation (activation or inhibition) and their relative likelihood. Only kinases with at least 5 phosphosite targets present in the analyzed phosphoproteomic data were considered for this analysis.

### Gene set overrepresentation analysis

Proteins with an adjusted p-value < 0.05 and a fold-change greater than 2 or smaller than 0.5, were defined as up-or downregulated proteins, respectively. Overrepresentation analysis of Reactome pathways in these lists of significant proteins was performed using the compareCluster function of the clusterProfiler R package (v. 4.10.1).

### Two-way ANOVA

Two-way ANOVA was performed on the filtered and normalized phosphoproteome data using limma (v. 3.58.1) with a design that includes the cell line (MCF7 WT, LTED or TAMR) and the time point of EGF stimulation (design = model.matrix(∼cell_line∼time)). This design was fir to the phosphoproteome data set using lmFit function and statistics was performed using the eBayes function. P-values were adjusted for multiple hypothesis testing with the Benjamini-Hochberg method.

Reactome gene set enrichment analysis was performed on the proteins that were found to be significantly differentially phosphorylated, using the enrichPathway function of the ReactomePA package (v. 1.46.0).

To identify overrepresented kinases that are responsible for the phosphorylation of these ANOVA-significant sites, hypergeometrical testing was performed using the phyper function of the stats R package (v. 4.3.1).

### Data visualization

Used functions that are not contained in base R: dotplot (clusterProfiler, v. 4.10.1), pheatmap (pheatmap, v. 1.0.12), textplot (wordcloud, v. 2.6), fviz_pca_ind (factoextra, v. 1.0.7).

For illustrations, Adobe Illustrator 2024 was used.

### Reporting summary

Further information on research design is available in the Nature Research Reporting Summary linked to this article.

## DATA AVAILABILITY

The mass spectrometry proteomics data have been deposited to the ProteomeXchange Consortium via the PRIDE^78^ partner repository with the dataset identifier PXD063393.

## CODE AVAILABILITY

All data files (normalized protein or phosphosite data sets, statistics outputs), as well as the code used for data quality control, normalization and creating the plots, are available on GitHub (https://github.com/schwarlu1691995/PAK1_manuscript).

## Notes

### Competing Interest Statement

The authors have declared no competing interest.

