## supplementary figures for "PAK1 activation drives divergent resistance mechanisms to aromatase inhibition and Tamoxifen in a luminal A breast cancer model"

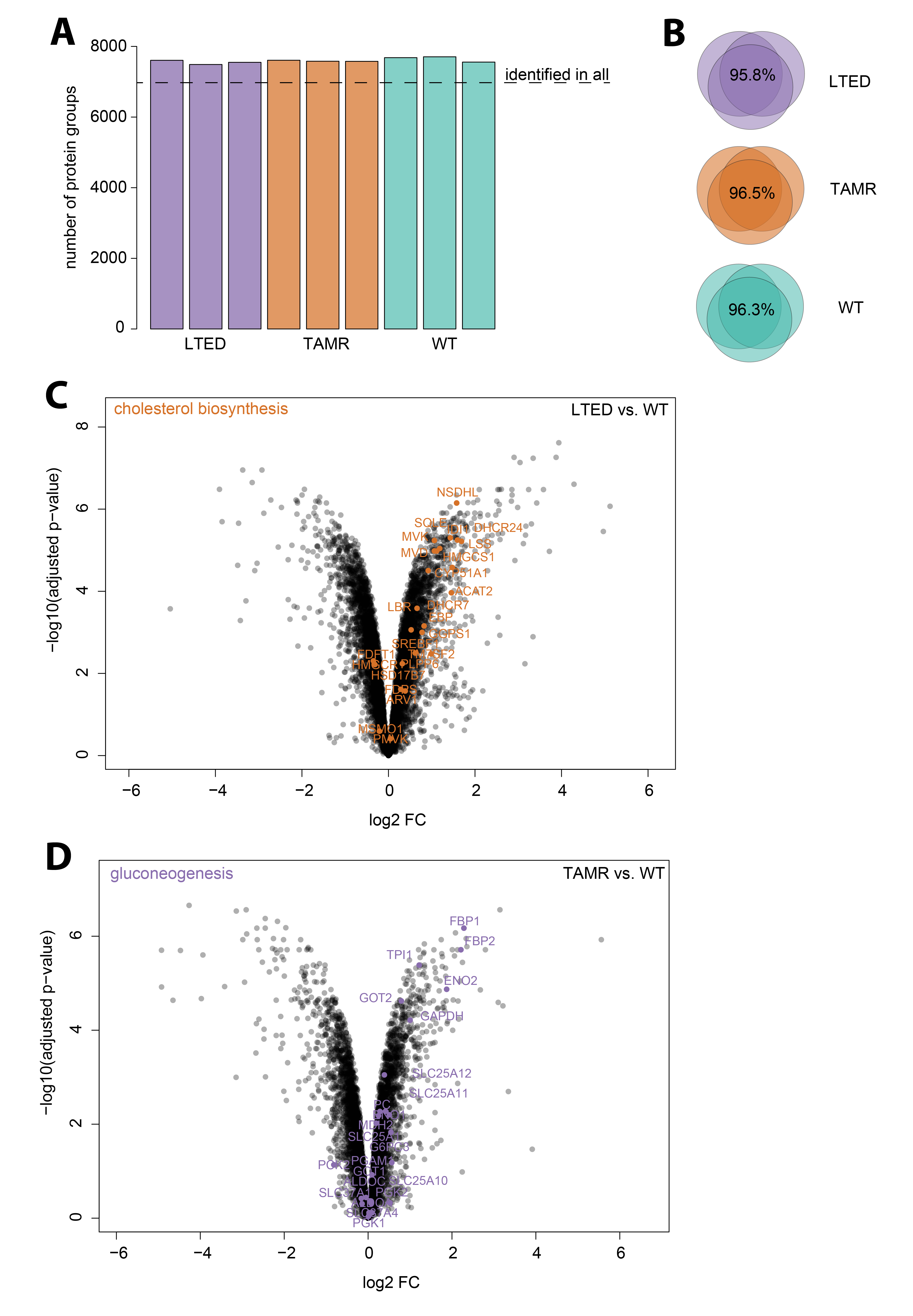

**Suppl. Figure 1: Full proteome adaptations of MCF7 LTED and TAMR.**

Cells were lysed in RIPA buffer, followed by 30 minutes of DNase/Benzonase treatment on ice. Protein clean-up and digestion was performed using manual SP3 with 500 µg protein input. Peptides were enriched for phosphorylation and both the full and phosphoproteome fractions were analyzed by LC-MS/MS for 120 minutes in DIA mode. Peptides and proteins were identified and quantified using Spectronaut (v17) in directDIA+ mode.

(A) Numbers of identified protein groups in the three biological replicates. (B) Overlap in identified protein groups between the biological replicates. (C, D) Statistical analysis was performed using the eBayes function of the limma R package. Reactome pathway overrepresentation analysis was performed, using a threshold of fold-change > 2 (UP) or < 0.5 (DOWN) and an adjusted p-value < 0.05. Log2 fold-changes and -log10 adjusted p-values are shown for all proteins compared to WT. Proteins involved in the significantly overrepresented pathways “cholesterol biosynthesis” and “gluconeogenesis” are highlighted in LTED (C) and TAMR (D), respectively.

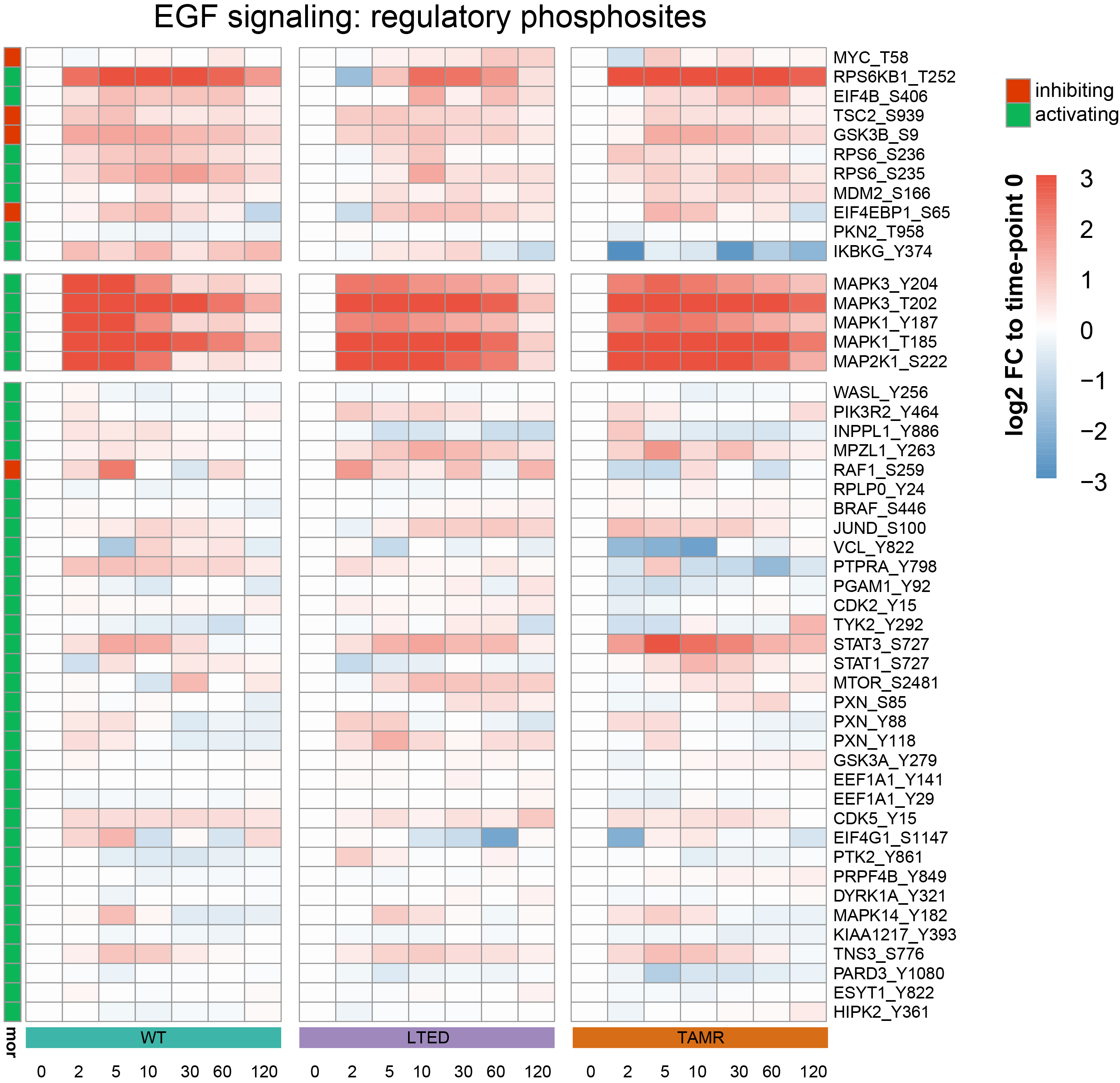

**Suppl. Figure 2: Known regulatory phosphosites of EGF signaling.**

(A) Time course of selected phosphosites involved in EGF signaling (according to PTMSigDB^68^) in response to EGF stimulation. The color represents the mean log2 fold-change to time point 0 (n = 3 biological replicates).

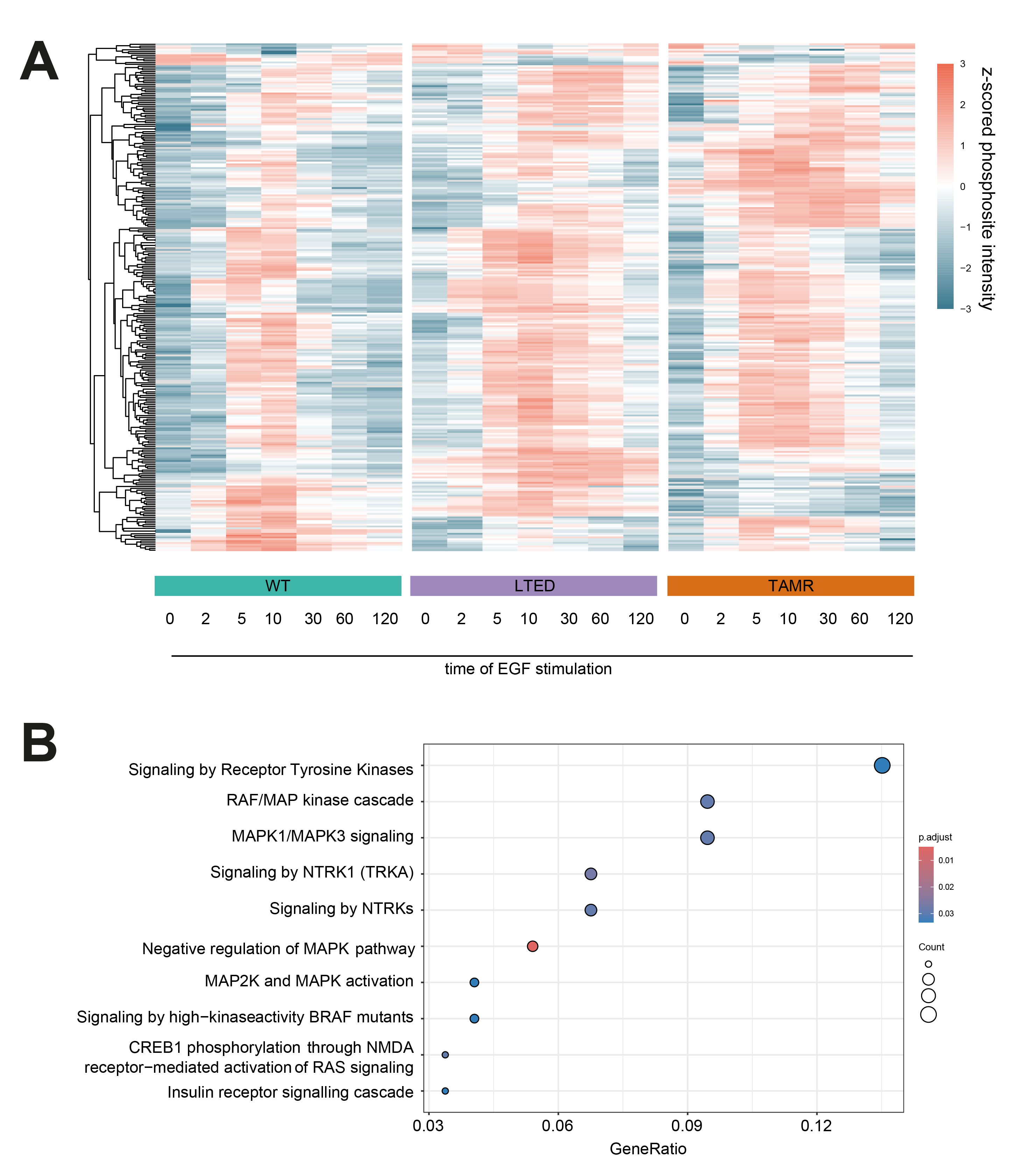

**Suppl. Figure 3: Two-way ANOVA analysis of phosphosites over time of EGF stimulation and between cell lines.**

(A) Z-scaled intensities of phosphosites found the be significantly different over time and between the cell lines in a 2-way ANOVA analysis. (B) Reactome gene set enrichment of the proteins phosphorylated in A.

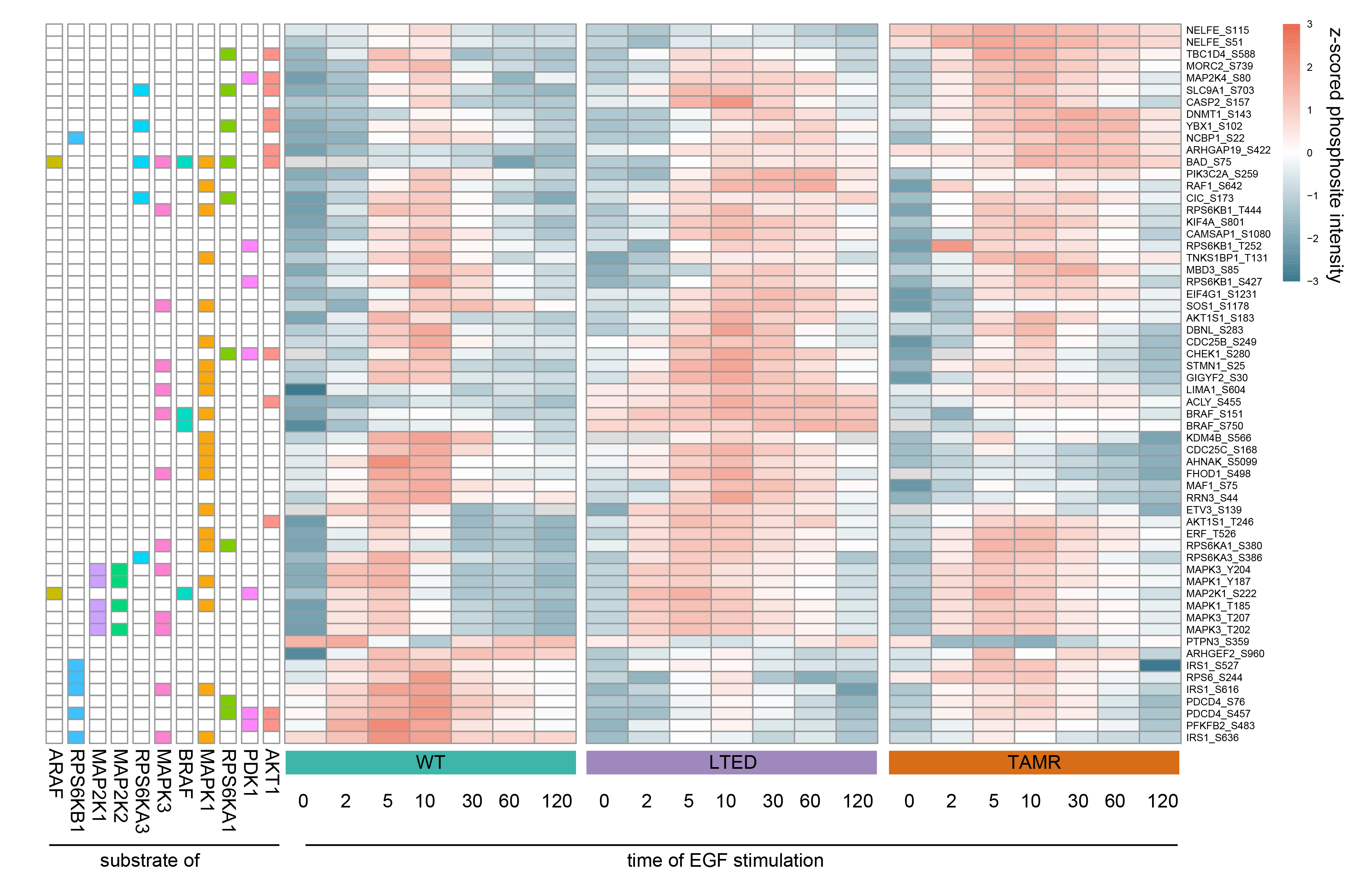

**Suppl. Figure 4: Kinase annotation of significant phosphosites from two-way ANOVA over time and cell lines.**

Phosphosites from Suppl. Figure 3 that have a responsible kinase annotated in Omnipath^30^. Overrepresentation analysis of kinases assigned to these sites revealed significant overrepresentation of the ten indicated kinases.

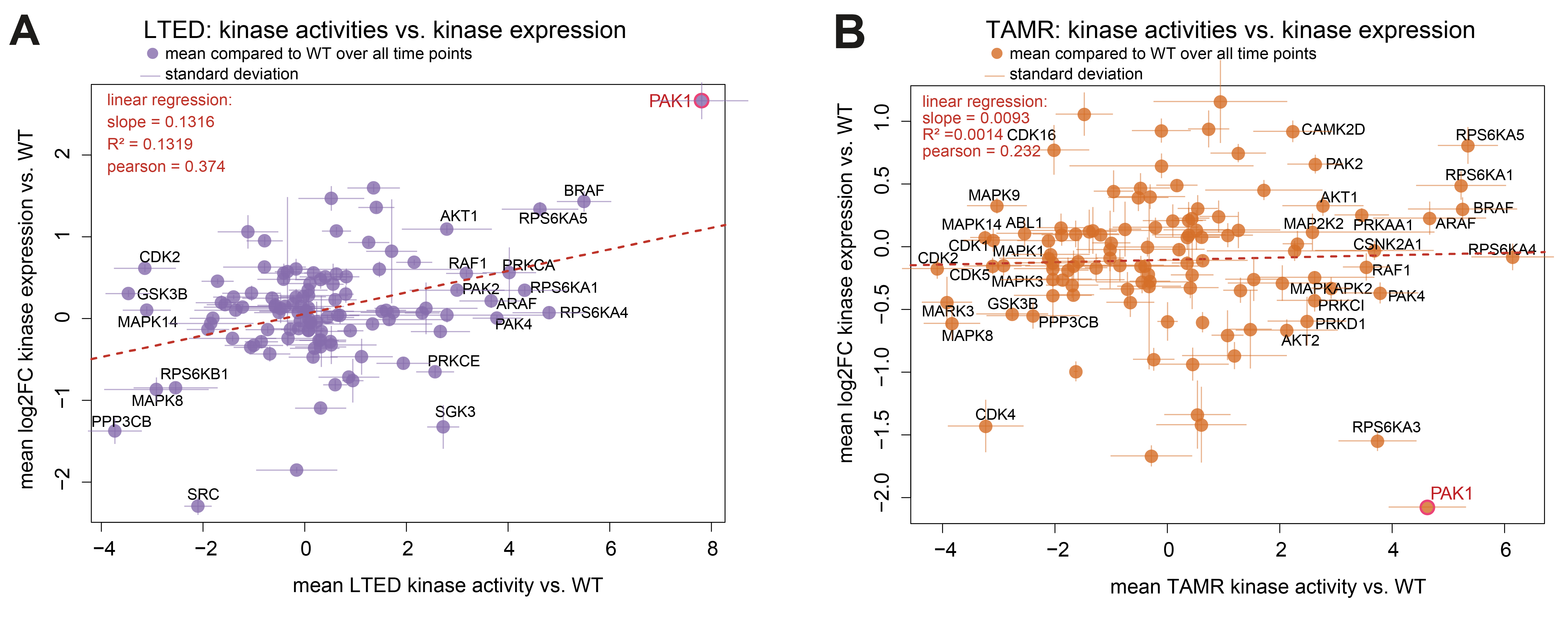

**Suppl. Figure 5: Comparison of kinase activities and expression levels.** (A, B) Mean kinase activity compared to WT at each time-point and mean kinase abundance levels as log2 fold-change compared to WT at each time point.

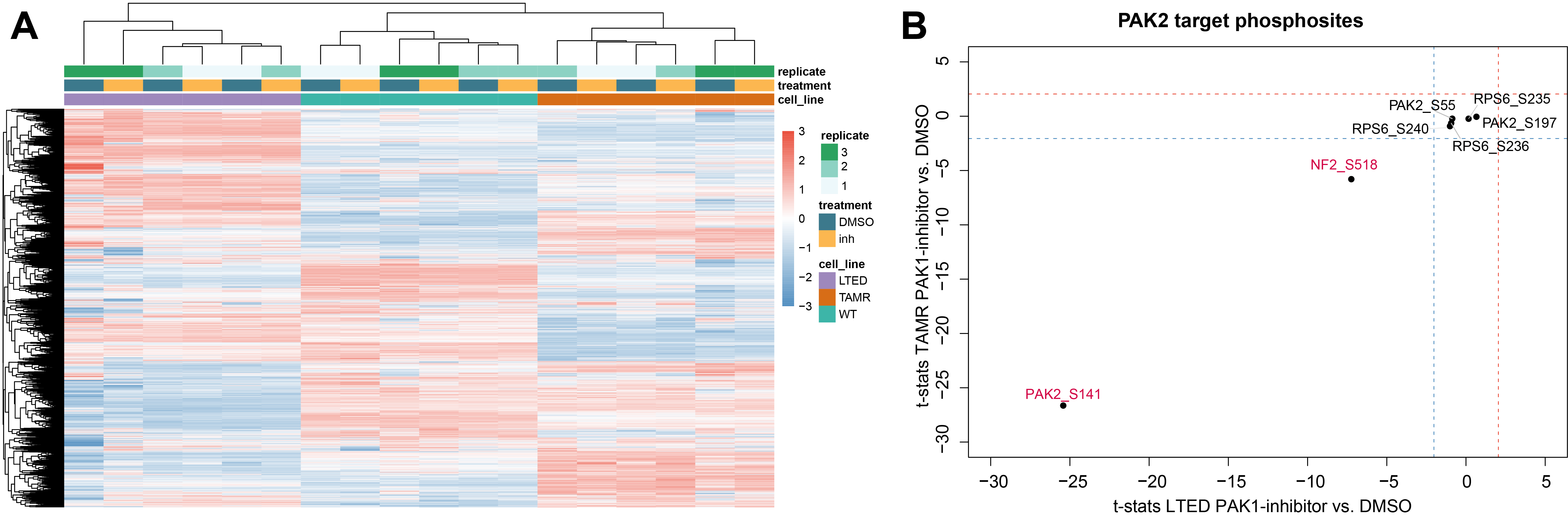

**Suppl. Figure 6: Proteome analysis following PAK1-inhibitor treatment.**

MCF7 cell lines (WT, LTED, TAMR) were treated with PAK1 inhibitor for 60 minutes, followed by LC-MS/MS analysis. (A) Z-scored heatmap and unsupervised hierarchical clustering of protein quantifications. No significant changes after 1h of PAK1 inhibition in any cell line. (B) T-stats of known PAK2 target sites (PAK1 inhibitor vs. DMSO) in LTED and TAMR.
